# Minimizing activation of overlying axons with epiretinal stimulation: The role of fiber orientation and electrode configuration

**DOI:** 10.1101/245266

**Authors:** Timothy Esler, Robert R. Kerr, Bahman Tahayori, David B. Grayden, Hamish Meffin, Anthony N. Burkitt

## Abstract

*Objective*. Currently, a challenge in electrical stimulation of the retina is to excite only the cells lying directly under the electrode in the ganglion cell layer, while avoiding excitation of the axons that pass over the surface of the retina in the nerve fiber layer. Since these passing fibers may originate from distant regions of the ganglion cell layer. Stimulation of both target retinal ganglion cells and overlying axons results in irregular visual percepts, significantly limiting perceptual efficacy. This research explores how differences in fiber orientation between the nerve fiber layer and ganglion cell layer leads to differences in the activation of the axon initial segment and axons of passage. *Approach*. Axons of passage of retinal ganglion cells in the nerve fiber layer are characterized by a narrow distribution of fiber orientations, causing highly anisotropic spread of applied current. In contrast, proximal axons in the ganglion cell layer have a wider distribution of orientations. A four-layer computational model of epiretinal extracellular stimulation that captures the effect of neurite orientation in anisotropic tissue has been developed using a modified version of the standard volume conductor model, known as the cellular composite model. Simulations are conducted to investigate the interaction of neural tissue orientation, stimulating electrode configuration, and stimulation pulse duration and amplitude. *Main results*. The dependence of fiber activation on the anisotropic nature of the nerve fiber layer is first established. Via a comprehensive search of key parameters, our model shows that the simultaneous stimulation with multiple electrodes aligned with the nerve fiber layer can be used to achieve selective activation of axon initial segments rather than passing fibers. This result can be achieved with only a slight increase in total stimulus current and modest increases in the spread of activation in the ganglion cell layer, and is shown to extend to the general case of arbitrary electrode array positioning and arbitrary target neural volume. *Significance*. These results elucidate a strategy for more targeted stimulation of retinal ganglion cells with experimentally-relevant multi-electrode geometries and readily achievable stimulation requirements.

## I. Introduction

There has been significant progress over the past decade in the development of retinal prostheses for sufferers of retinal pathologies such as retinitis pigmentosa. Clinical trials of retinal prostheses have found that patients can reliably report visual percepts arising from stimulation, and can perform simple identification tasks [1–8]. Although progress to date is highly encouraging, many aspects of the performance of retinal prostheses remain limited, hinging on the ability of these devices to target either specific retinal cell types [9, 10] or more precise retinal volumes [1, 3, 11]. In the case of epiretinal stimulation, a factor limiting performance is the inability of electrical stimulation to preferentially activate target neuronal structures in the ganglion cell layer (GCL) while avoiding activation of overlying axons in the nerve fiber layer (NFL) [1, 11–19], illustrated in Fig. 1.

**Fig. 1.**
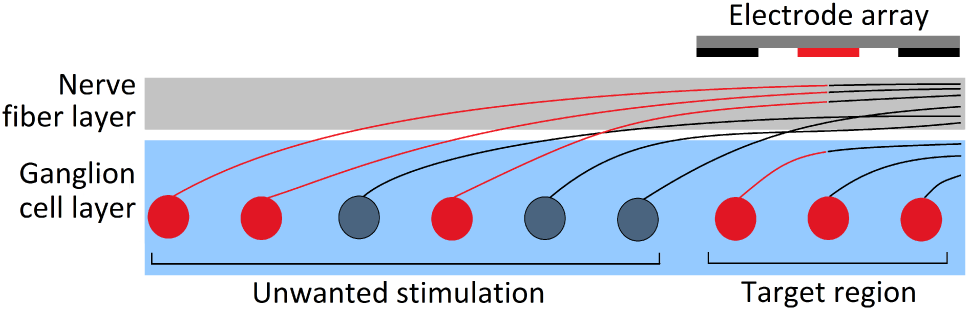
Unwanted stimulation of retinal ganglion cell axons of passage in the nerve fiber layer that pass close to stimulating electrodes. Activated retinal ganglion cells are colored red.

The two-dimensional organization of cells in the retina with respect to incident light or electrical stimulation is lost as the axons of retinal ganglion cells (RGCs) traverse the inner surface of the retina in the NFL. As a result of this structure, epiretinal electrical stimulation faces the challenge of stimulating the deeper, favourably-organized GCL while minimizing activation of axons of passage (AOPs) in the NFL. Irregular visual percept shapes are commonly described by recipients of epiretinal implants due to stimulation of axons of passage [13, 15, 20, 21]. This effect has been confirmed experimentally and in simulations, and results in a reduction in the spatial selectivity of epiretinal stimulation [11, 15–17, 20–22].

A potential way to minimise activation of axons of passage in the retina is to take advantage of differences in neurite orientation in the NFL and GCL. The direction of overlying passing axon tracts represents the dominant fiber orientation in a given location in the NFL. These axons are packed together as mostly parallel fibers [11, 13]. As a result, current flow from epiretinal electrical stimulation spreads through retinal tissue in a highly anisotropic way. In contrast to the distal RGC axons in the NFL, proximal axon regions such as the axon initial segment (AIS), located in the GCL, have a much wider distribution of orientations as they pass out from the soma. It is expected that, based on these anisotropic tissue characteristics, the orientation of a neurite in retinal tissue can have a significant effect on its activation. However, a common approximation employed by existing computational models of epiretinal stimulation is that the retinal layers are isotropic [11, 16, 18, 23]. In order to assess the effect of neurite orientation, and its interaction with different multi-electrode configurations, computational models of current flow and axonal activation should be developed that can describe the anisotropic characteristics of different retinal layers.

In the absence of detailed data on the anisotropy of the NFL, an alternative approach is to derive layer anisotropy from first principles using a geometric description of the axonal units that comprise the tissue. The cellular composite model, introduced by Meffin et al. [24–27] provides a modeling framework that accomplishes this while addressing a number of limitations of conventional volume conductor models. In order to more accurately capture the structural and temporal properties of neural tissue and to guarantee model self-consistency, the cellular composite model maps extracellular current to voltage using an expression for impedance derived directly from the geometry and physiology of the NFLs microscopic constituent axons.

In addition to intrinsic tissue anisotropy, RGC activation will also depend on the orientation of the applied electric field. One existing modeling study by Rattay and Resatz [11] assessed the influence of electric field orientation with respect to neurites in the NFL. This study showed that, by orientating long, rectangular electrodes parallel to axons in the NFL, the activation of those axons could be reduced. The basis for this result is that the membrane potential response of an axon to extracellular stimulation is approximately proportional to the activating function: the second spatial derivative of the induced extracellular potential along the axon’s length [28]. By *flattening* the extracellular potential along the length of the axon using long parallel electrodes, the activation of the axon is minimised.

The aim of this paper is to demonstrate a multi-electrode stimulation strategy for the avoidance of activation of axons of passage, while achieving focal activation of axon initial segments the in GCL. We present a model that captures the effect of both electric field orientation introduced by multi-electrode stimulation, and the effect of the highly anisotropic geometry of the NFL. Simulation results are presented that illustrate the achievable levels of preferential activation for one-, two-, and four-electrode configurations. An exploration of the effects of electrode-retina separation distance and pulse duration are presented, as well as the effects of different strategies on key performance metrics: required stimulus current, GCL activation, and activation radius. The suggested multi-electrode array strategy will then be validated against a more general set of electrode geometries and target volumes.

## II. Methods

### A. Distribution of orientations in the ganglion cell layer

To quantify the distribution of orientations of proximal axons in the GCL, we analysed 777 RGC reconstructions obtained from NeuroMorpho.org [29–40]. Cell morphologies were imported and processed in MATLAB (The Mathworks, Release 2016a) with the assistance of the third party TREES toolbox [41]. For each cell for which enough of the axon was reconstructed, the difference in the orientation between the AIS (defined as the segment from 40*μ*m to 80*μ*m from the soma [42]) and various locations along the axon was calculated, as illustrated in Fig. 2(a). Fig. 2(b) and 2(c) show the proportion of cells with orientation differences in different ranges. As shown in Fig. 2(b), the orientation difference in the *x-y* plane approaches a uniform distribution for locations 500*μ*m (or more) distal from the AIS. In contrast, Fig. 2(c) shows that there is very little change in orientation between the AIS and AOP in terms of altitudinal orientation, indicating that axons remain predominantly parallel to the surface of the retina along their length. Based on the knowledge that fibers in the NFL are approximately parallel at a given location, this analysis suggests an approximately circular (but not spherical) uniform distribution for AISs in the GCL, and this distribution will be used in this paper.

**Fig. 2.**
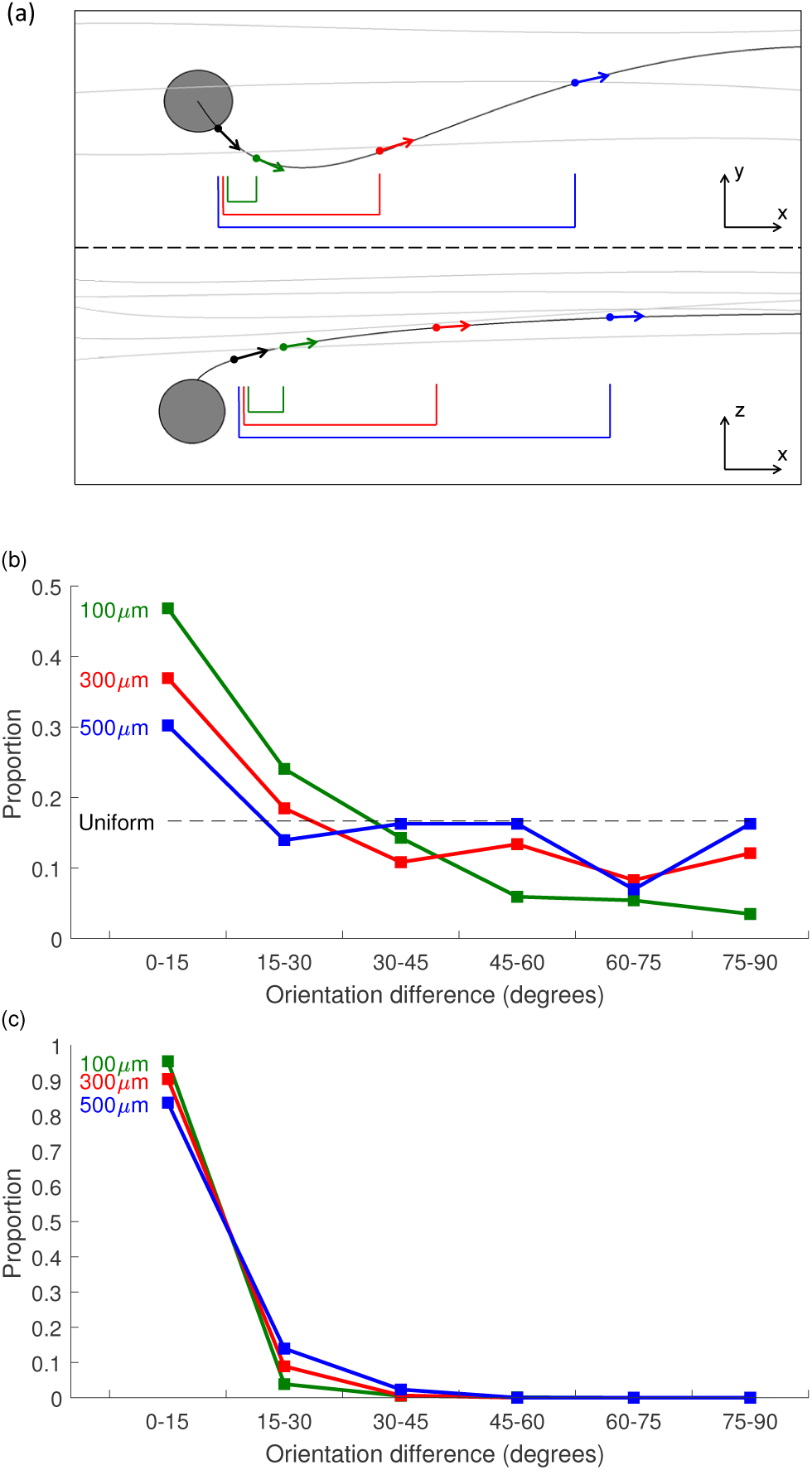
Change in axon orientation relative to the orientation of the AIS at 100*μ*m (green), 300*μ*m (red) and 500*μ*m (blue) along the axon from the soma. The (b) azimuthal (i.e. *x-y*) and (c) altitudinal (i.e. *z*) change in orientation between the AIS and various locations along the axon are shown separately, as illustrated in (a). Distributions were calculated from all available RGC reconstructions on NeuroMorpho.org. Due to variation in the length of axon reconstructions, each trace is calculated using a different number of cells (100*μ*m - 777 cells, 300*μ*m - 157 cells, 500*μ*m - 42 cells).

### B. Tissue geometry and governing equations

The model employed here uses a two-stage volume conductor framework. The first stage models the field of extracellular electric stimulation due to the stimulating electrodes. The second stage uses the calculated extracellular potential or current from the first stage as input into a passive neurite model to calculate membrane potential. The present modeling approach uses a four-layer description of retinal geometry for stage one (Fig. 3). The modeled layers are the insulating substrate of the electrode array, the vitreous, the nerve fiber layer, and a lumped approximation of the remaining retinal layers, including the ganglion cell layer. The conductivity/admittivity and directional dependence properties of each layer are presented in Table I. Admittivity is a spatially- and temporally-dependent generalization of conductivity and is this inverse of impedivity, meaning that it contains both resistive (real) and reactive (imaginary) parts. The anisotropic admittivity of the NFL is incorporated into the complex admittivity kernel provided by the cellular composite model of Meffin et al. [26].

**Fig. 3.**
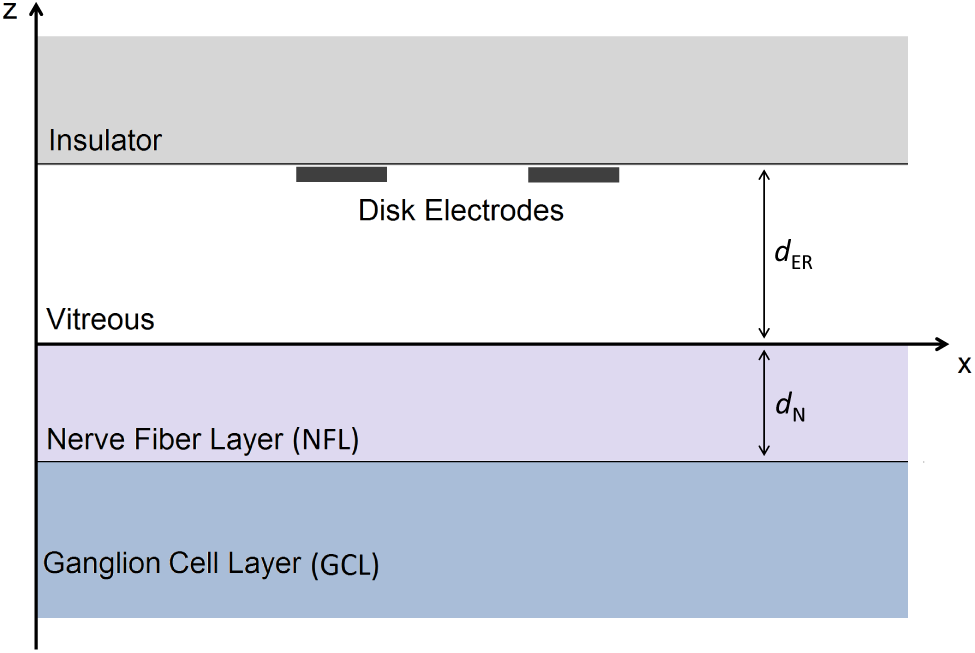
Geometry of the four layer model of the retina. Modeled layers include the insulator, vitreous, NFL, and GCL. The insulator is assumed to have zero conductivity and is modeled using a zero flux boundary condition. The GCL is assumed to have infinite extent in the z-direction. The distance from electrodes to the retinal surface and the thickness of the NFL are denoted by *d*_ER_ and *d*_N_, respectively.

**TABLE I.** Conductivity and thickness of modeled layers

| Layer | Directional dependence | Conductivity (S/m) | Thickness ($\mu\text{m}$) |
| --- | --- | --- | --- |
| Insulator (I) | Isotropic | 0 | Infinite extent |
| Vitreous (V) | Isotropic | 1.78 | 10-500 (based on location of electrode array) |
| NFL (N) | Anisotropic | see Equations (13)-(14) | 100 [43–46] |
| GCL (G) | Isotropic | 0.1 | Infinite extent |

Due to the (approximately) zero conductivity property of the insulator layer, the effect of this layer is included via a zero current boundary condition applied at the insulator-vitreous layer boundary. Electrodes are modeled as flat, circular disks lying on this boundary.

The flow of current in the extracellular space in each layer is described by a separate Poisson-type equation, allowing for differing tissue conductivities in each layer, with the current delivered by disk electrodes entering as an explicit term on the right-hand-side of the vitreous layer continuity equation:

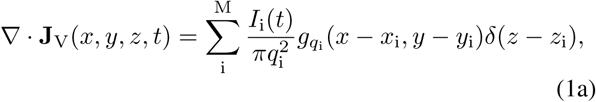

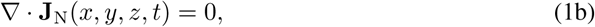

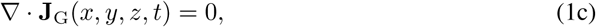

where **J***_α_* is the extracellular current density in layer *α* and each set of (*x_i_*, *y_i_*, *z_i_*) represents the three-dimensional location of each of the M electrodes. Each electrode has a radius, *q_i_*, and stimulus current waveform, *I_i_*(*t*). The function 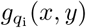 is the unit circular step function of radius *q_i_* in the *x-y* plane and *δ*(*z*) represents the Dirac delta function. For this model, each layer boundary is approximated by an infinite flat plane parallel to the *x-y* plane, so that the set of electrode heights, *z_i_*, are equal. Furthermore, if the origin is fixed on the vitreous-NFL boundary, then *z_i_* is equivalent to the electrode-retina separation distance, *d*_ER_.

A generalized form of Ohm’s Law is used to describe extracellular current density, which is governed by each layer’s admittivity kernel. This admittivity kernel incorporates the dependence of the extracellular current density on the electric field at previous times and at remote locations in the extracellular space. These atypical dependencies arise due to the passage of current across the cellular membrane and through the intracellular space. The relationship between extracellular potential and current density is described by

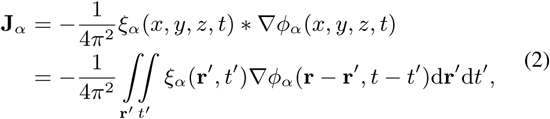

where *ξ_α_* is the 3×3 admittivity kernel and *ϕ_α_* is the extracellular potential of layer *α* ∈ {V, N, G}. ∇ = [*∂/∂x; ∂/∂y; ∂*/*∂*z] is the differential operator. In the most general case, where *ξ_α_* varies in three spatial dimensions and time, * represents a convolution over three spatial dimensions and time. For brevity, the spatial coordinates (*x, y, z*) have been represented by the vector **r** in the integral expression for the convolution.

Expanding Equation (2) for each layer gives

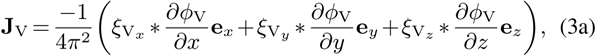

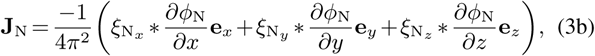

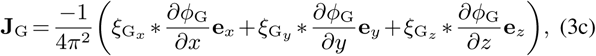

where *e_x_*, *e_y_*, and *e_z_* are unit vectors in *x, y*, and *z* directions, respectively. Note that each of the dimension-specific admittivity terms in the above expression may have (*x*, *y*, *z*, *t*) dependence.

By assuming that within each layer tissue admittivity is independent of *z*, we can reduce the above four dimensional convolutions to three dimensions. Using the *x*-component of the admittivity in the vitreous layer to illustrate, we have

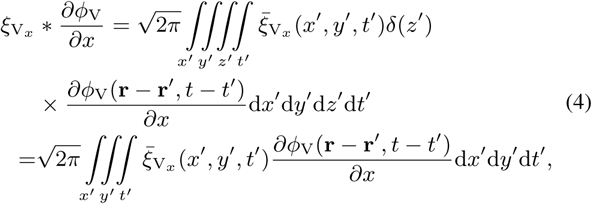

where 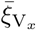 is the *x*-component of the admittivity in the vitreous layer at a given point in (*x, y*, *t*) space, which does not vary with *z* within a layer. Identical simplifications can be shown for each of the nine convolutions in Equations (3). Subsequently, this will allow for the removal of these convolutions using a Fourier transform in three dimensions instead of four.

For layers with infinite extent in the *x*- and *y*-directions, as in the present model, boundary conditions are specified at the layer boundaries:

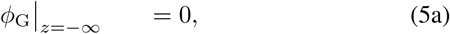

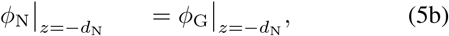

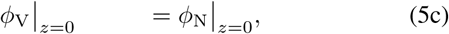

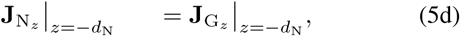

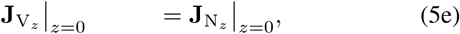

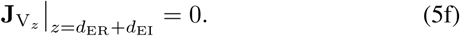

These boundary conditions ensure the described system has finite energy (Equation (5a)), current density and voltage vary continuously across layer boundaries (Equations (5b-e)), and no current can flow into the insulating substrate (Equation (5f)).

Since the current sources are at the same *z*-location as the insulator’s zero current condition, we initially define the geometry such that the insulator is separated from the electrodes by some distance, *d*_EI_. This was eliminated subsequently by computing the limit from above as *d*_EI_ goes to zero.

Solution of the system of elliptic partial differential equations defined by Equations (1) and (3) using layer boundary conditions (5) yields expressions for the extracellular potential in each layer. In order to find an analytic solution to this system, Fourier domain approaches are applied to reduce the convolutions shown in Equation (3) to multiplications. All Fourier domain transformations performed in these analyses are of the following form:

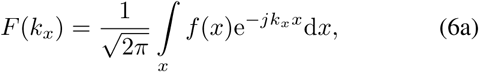

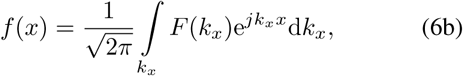

where *k_x_* and *F*(*k_x_*) are the Fourier transform pairs of *x* and *f* (*x*), respectively.

### C. Solution of volume equations

Taking the Fourier transform of each of Equations (1) with respect to *x, y*, and *t* gives the following set of equations:

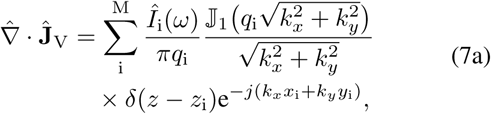

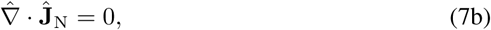

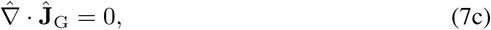

where the hat symbol (ˆ) indicates the Fourier transform of the specified quantity with respect to *x, y*, and *t*, with Fourier transform pairs *k_x_*, *k_y_*, and *ω*, respectively. 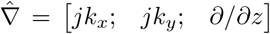 is the Fourier transform of the differential operator and 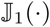 represents the Bessel function of the first kind of order 1. Similarly, taking the Fourier transform of Equations (3) yields

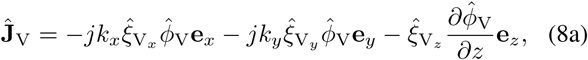

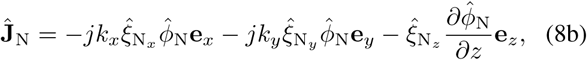

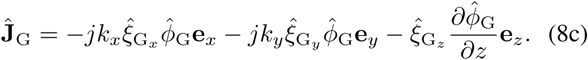

Substituting Equations (8) into (7), the system may be written as

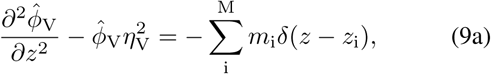

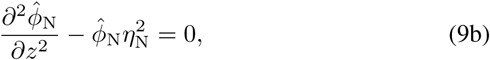

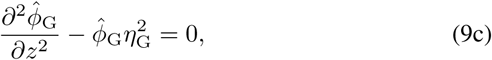

where

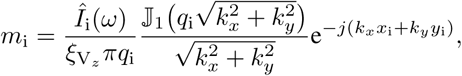

and

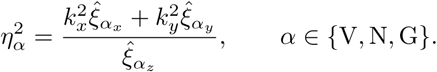

Solutions to Equations (9) are of the form

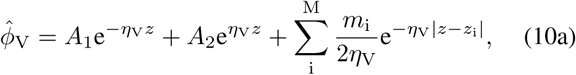

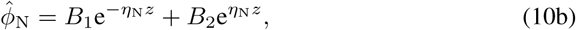

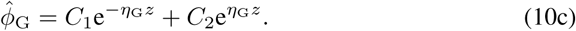

By substituting the boundary conditions in Equations (5) into Equations (8) and (10), the following simultaneous equations define the system’s constants of integration

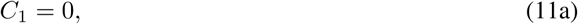

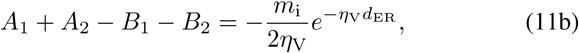

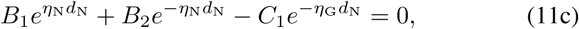

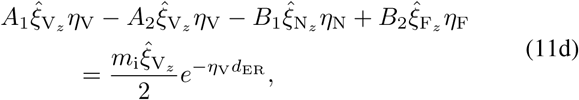

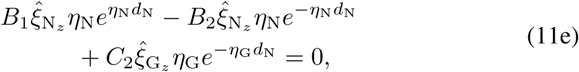

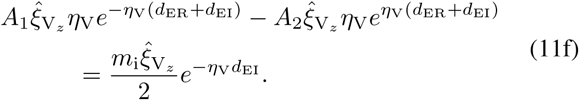

Given the complexity of the resultant expressions, this set of equations was solved with the symbolic mathematics engine, Mathematica (Wolfram Research, Version 10). Note that the integration constants are obtained separately for each electrode. Owing to the model’s linearity, multi-electrode simulations are implemented via the superposition of the electric field generated from several single-electrode simulations. In order to eliminate *d*_EI_ from the resulting solution, the right limit as *d*_EI_ goes to zero was also computed with Mathematica.

### D. Admittivity of the nerve fiber layer

As shown in Table I, the insulator, vitreous, and ganglion cell layers are described using scalar conductivities. This is equivalent to these layers having a spatially- and temporally-independent admittivity term that is also isotropic. Like the NFL, the GCL consists of active neural tissue, and hence would be expected to have some capacitive response to stimulation, which would lead to a nonzero permittivity. However, since the GCL is predominantly composed of cell somas, the volume proportion taken up by neural membrane is much lower than that of the NFL, which is predominantly composed of densely packed and thinner fibers. For this reason, the GCL, as for the insulator and vitreous layers, is assumed to have a constant conductance, and is considered to be isotropic:

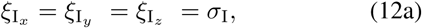

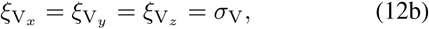

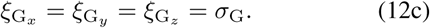

The cellular composite model provides expressions for both the tissue admittivity kernel for the NFL and for the membrane potential of a neurite given the extracellular potential along its axis (irrespective of which layer the neurite is in) [26].

We assume that in the NFL fibers are oriented in the *y*-direction. Then, in the time and space domains, the NFL admittivity is given by

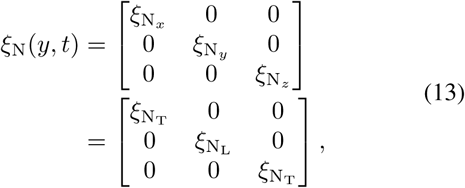

where

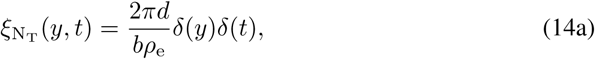

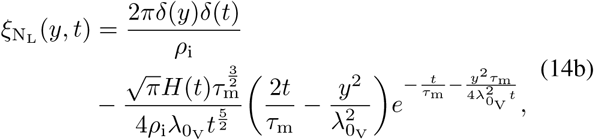

in which 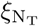 and 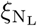 are the transverse and longitudinal components of the admittivity kernel, respectively, and *H* is the Heaviside step function. The remaining terms, *d*, *b*, *ρ*_e_, *ρ*_i_, *τ*_m_, and 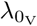 each represent different physical or electrical properties of the tissue and are defined in Table II. The above expressions reduce to the following by taking the Fourier transform with respect to *y* and *t*:

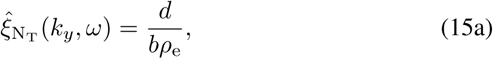

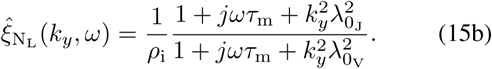

**TABLE II.** Model parameters

| Parameter | Description | Unit |
| --- | --- | --- |
| $a$ | Neurite radius | m |
| $b$ | Outer cylinder radius | m |
| $d$ | Width of extracellular sheath, $d = b - a$ | m |
| $\rho_i$ | Intracellular resistivity | $\Omega \text{ m}$ |
| $\rho_e$ | Extracellular resistivity | $\Omega \text{ m}$ |
| $r_i$ | Intracellular resistance per unit length, $r_i = \rho_i / (\pi a^2)$ | $\Omega / \text{m}$ |
| $r_e$ | Extracellular resistance per unit length, $r_e = \rho_e / (\pi(b^2 - a^2))$ | $\Omega / \text{m}$ |
| $R_m$ | Membrane unit area resistance | $\Omega \text{ m}^2$ |
| $r_m$ | Membrane unit length resistance, $r_m = R_m / (2\pi a)$ | $\Omega \text{ m}$ |
| $C_m$ | Membrane capacitance per unit area | $\text{F} / \text{m}^2$ |
| $\tau_m$ | Membrane time constant, $\tau_m = R_m C_m$ | s |
| $\lambda_{0J}$ | Static electrotonic length constant for current density boundary conditions, $\lambda_{0J} = \sqrt{r_m / (r_e + r_i)}$ | m |
| $\lambda_J(\omega)$ | Frequency-dependent electrotonic length constant for current density boundary conditions, $\lambda_J(\omega) = \lambda_{0J} / \sqrt{1 + j\omega\tau_m}$ | m |
| $\lambda_{0V}$ | Static electrotonic length constant for voltage boundary conditions, $\lambda_{0V} = \sqrt{r_m / r_i}$ | m |
| $\lambda_V(\omega)$ | Frequency-dependent electrotonic length constant for voltage boundary conditions, $\lambda_V(\omega) = \lambda_{0V} / \sqrt{1 + j\omega\tau_m}$ | m |
| $d_N$ | Nerve fiber layer thickness | m |
| $d_{ER}$ | Electrode-retina separation distance | m |
| $d_{EI}$ | Electrode-insulator separation distance, set to zero in final solution | m |
| $q_i$ | Radius of disc electrode $i$ | m |
| $x_i, y_i, z_i$ | Location of center of electrode $i$ | m |

Specific layer admittivities and conductivities define the level of anisotropy for each layer as well as unique spatiotemporal dependencies. For the NFL and GCL, the form of the admittivity represents the assumed distribution of fiber orientations in each layer. The NFL is modeled as a parallel fiber bundle (anisotropic) and fibers in the GCL are modeled as having a uniform distribution of orientations (isotropic). Equations (14) describe a non-local, non-instantaneous admittivity for the NFL, which is derived from an accurate characterization of the spatiotemporal electrical properties of individual neurites that comprise the tissue [26, 47].

### E. Neurite equations

Stage two of the cellular composite model involves the calculation of the passive membrane potential in the neurite of interest, in either the NFL or the GCL. This is achieved using the neurite equations of Meffin et al. [24], which provide expressions for membrane activation due to modes of current flow that are both longitudinal and transverse with respect to the fibers. Expressions for the membrane potential along a single fiber in a fiber-bundle with orientation parallel to the y-axis (as in the NFL) are supplied in the *x,y,t*-Fourier domain by the cellular composite model:

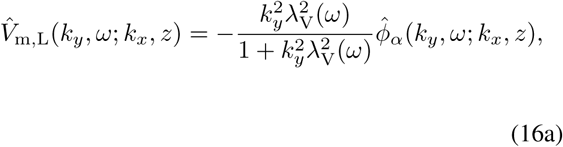

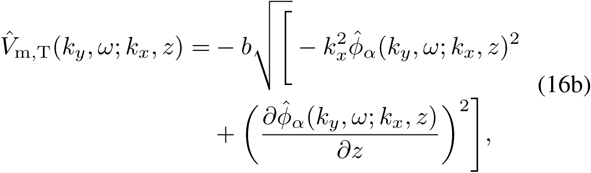

where 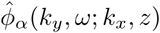 is the extracellular potential along the neurite axis for a straight neurite oriented parallel to the *y*-axis at a point (*k_x_, z*). Equation (16a) is a Fourier domain representation of the cable equation for extracellular stimulation, and indicates the dependence of *V*_m,L_ on the second spatial derivative of the extracellular potential in the direction of the neurite, known as the activating function [28]. Here, the activating function is represented in the Fourier domain as 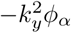.

Extension of expressions for the longitudinal and transverse components of the membrane potential to straight neurites of arbitrary *x-y* orientation allows for analysis of fibers in both the NFL and GCL.

### F Generalisation of neurite equations

Owing to the high anisotropy of the NFL, fibers with different orientations, whether they are in the NFL or lower layers, will achieve different levels of activation given an applied stimulus. The volume conductor model derived above assumes that membrane potential is calculated for fibers with a single orientation: parallel to the NFL fiber bundle.

The model includes distinct stages for the calculation of extracellular potential (which captures the anisotropy of the NFL) and the calculation of membrane potential. As a result, it is possible to calculate the membrane potential for an unbranched axon with arbitrary morphology if we assume that the orientation of the axon of interest has a negligible effect on the distribution of extracellular potential as a whole. We can first calculate extracellular potential in the region of interest using Equations (10) and then sample it along the desired axon trajectory. This sampled extracellular potential can then be substituted into Equations (16) to yield the membrane potential along the axon.

Although this approach achieves a high level of generality, it requires an extra two computationally expensive Fourier transform calculations: one to convert the extracellular potential into the spatial domain from the spatial frequency domain to allow for sampling and another to convert the sampled data back into the frequency domain.

In the case of simple *x-y* rotations of straight axons, as assumed in the GCL, the required rotation can be performed directly in the Fourier domain, since rotation is preserved under unitary transformations such as the Fourier transform. Extracellular potential is thus calculated using a straightforward modification of Equations (10) in order to rotate the *k_y_*-axis (and equivalently the *y*-axis to align with the desired fiber orientation:

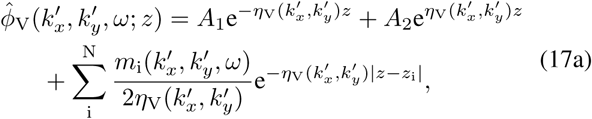

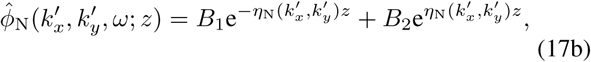

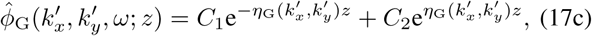

where, for a rotation in the *x-y* plane of *θ*, we use

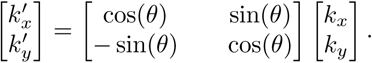

The corresponding rotated extracellular potential can then be applied directly to Equations (16).

### G. Simulation Methods

Simulation of a wide range of electrode and current waveform variations was conducted in MATLAB. All computations of induced extracellular potential and membrane potential were first calculated in the spatial and temporal frequency domains using Fourier domain solutions to the modelled system. The frequency representation of the longitudinal and first transverse components of a neurite's or a volume's membrane potential are summed together prior to calculating the inverse Fourier transform, yielding the final membrane potential.

The solution to the system described above is found in the Fourier domain with respect to the x and y spatial dimensions and the temporal dimension. Due to this, each simulation requires the calculation of extracellular and membrane potential in an entire spatial plane and for the full temporal extent of the simulation before the inverse Fourier transform is calculated. This process requires considerable memory resources and is facilitated using a custom parallel external memory (PEM) algorithm written in MATLAB.

This analysis considers only direct cell responses and neglects the effects of retinal networks. As such, the output of the passive membrane potential model is compared to pre-calculated membrane thresholds for the AIS and AoP to determine corresponding levels of activation. Pre-calculated thresholds are derived from experimental data using the method presented in Section III-A.

To determine the proportion of fibers activated at a given location within the retina, the activity of fibers with an appropriate range of orientations in the *x-y* plane is first calculated and then combined in a weighted sum, where the weights are sampled from an assigned distribution of orientations. For locations in the NFL, a single parallel orientation is assumed, whereas, for the GCL, a uniform distribution of orientations is applied in the *x-y* plane, as validated in Fig. 2.

We will describe and analyse the results of simulations of straight cylindrical neurites embedded in the modelled four-layer retinal structure. For all simulations, 100*μ*m diameter disk electrodes were used unless stated otherwise. For simulations of multi-electrode stimulation, electrodes were arranged in a regular grid with 200*μ*m center-to-center spacing between electrodes. Unless otherwise stated, stimuli used were cathodic-first, biphasic pulses with a pulse width of 200*μ*s. In addition, for all multi-electrode simulations, equal currents were applied to each electrode. All relevant parameter values used are presented in Table III. To compare the activation of AoPs in the NFL and AISs in the GCL, we consider characteristic axons located just (*z* = 10*μ*m) below the surface of their respective retinal layer, as structures at these locations are most sensitive to epiretinal stimulation.

**TABLE III.** Model parameter values

| Parameter | Value | Reference |
| --- | --- | --- |
| $a$ | 0.47 $\mu\text{m}$ | [29–40] |
| $d$ | 30 nm | [48] |
| $\rho_i$ | 0.7 $\Omega\text{m}$ | [49, 50] |
| $\rho_e$ | 0.7 $\Omega\text{m}$ | [49, 50] |
| $R_m$ | 1 $\Omega \text{ m}^2$ | |
| $C_m$ | 0.01 F/m<sup>2</sup> | |
| $d_N$ | 100 $\mu\text{m}$, unless stated otherwise | [43–46] |
| $d_{ER}$ | 10-500 $\mu\text{m}$ | |
| $q_i$ | 50 $\mu\text{m}$, unless stated otherwise | |

## III. Results

### A. Calculating membrane potential thresholds

Threshold potential values for the AIS and AOP have been determined from simulations that replicate the experimental procedures of Fried et al. [42]. These experiments found that a high-density sodium channel band was located at the RGC AIS (approximately 40–80*μ*m from the soma), resulting in a region of low stimulus threshold under epiretinal stimulation. A markedly higher stimulus threshold was observed at axonal locations either side of the AIS. The stimulus threshold was also shown to be decreased in the distal axon (or AOP), likely due to the axon moving up into the NFL where it is closer to the stimulating electrode. By matching the experimental electrode geometry, electrode location, neurite orientation, nerve fiber layer thickness, and stimulation frequency, Fried's experimentally-determined threshold stimulus currents were mapped to corresponding threshold membrane potentials in the computational model presented here.

In order to design simulations that most closely match the experimental methodology, nerve fiber layer thickness, *d*_N_, was set to 25*μ*m, appropriate for a rabbit retina. A single electrode with a radius, *q*, of 15*μ*m was used to deliver a single cathodic-first, biphasic pulse from a location 25*μ*m from the surface of the retina (*d*_ER_ = 25*μm*). Stimulus pulse amplitudes were chosen to approximately match the stimulus threshold levels reported by Fried et al., read from Figure 6A in [42]. Experimentally reported stimulus current thresholds for the AIS and the distal axon were then used as pulse amplitudes in simulations, which are illustrated in Fig. 4. These values happened to be approximately 20*μ*A for both the initial and distal axon due to the trade-off between proximity to the electrode array and the membrane threshold. The distal axon is closer to the stimulating electrode but has a higher membrane potential, whereas the AIS is further from the electrode but has a lower membrane threshold, resulting in a similar threshold stimulus current for each location. In this analysis, the AIS was assumed to be 5*μ*m below the surface of the GCL and the AOP was assumed to be centered in the NFL, 12.5*μ*m below the retinal surface. The experimental procedure of Fried et al. used narrow conical electrodes with no backing insulator and so the insulator layer was removed in these simulations. using these parameters, the maximum simulated membrane depolarisation achieved in an axon below the electrode corresponded to the relevant membrane threshold. Membrane thresholds were 12.09mV and 6.30mV for the AOP and the AIS, respectively.

**Fig. 4.**
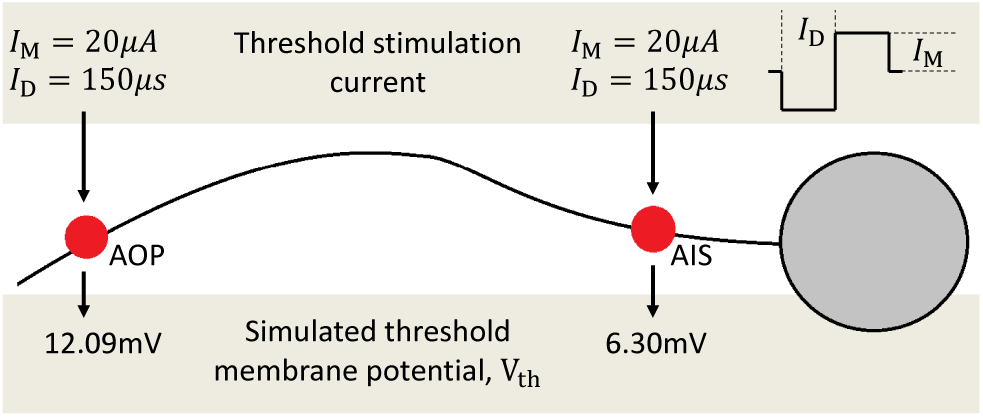
Simulation of experiments from Fried et al. [42], showing the current waveforms used for stimulation and the simulated membrane potential response. The maximum simulated membrane potential for each simulation corresponds to the the membrane threshold, *V*_th_, for that location in the axon.

### B. Comparison of one-, two-, and four-electrode configurations

To establish a basis for fiber orientation-dependent activation in the retina, simulations were run to determine the activation for a single characteristic AOP and AISs with a range of sampled azimuthal orientations. The geometry of these simulations is represented in Fig. 5(a). Figs. 5(b), (c), and (d) show the membrane potential resulting from stimulation with one, two, and four electrodes, respectively, for fibers with orientations illustrated in Fig. 5(a). Fig. 5(b) highlights the influence of the NFL anisotropy on the activation of GCL fibers of different orientations, with fibers orientated perpendicularly to the AOP experiencing 1.9 times the depolarisation of parallel fibers. Fig. 5(b) also highlights the problem being addressed by this research: although the target AISs in the GCL have a much lower membrane threshold, the proximity of the NFL to stimulating electrodes results in the preferential activation of AOPs.

**Fig. 5.**
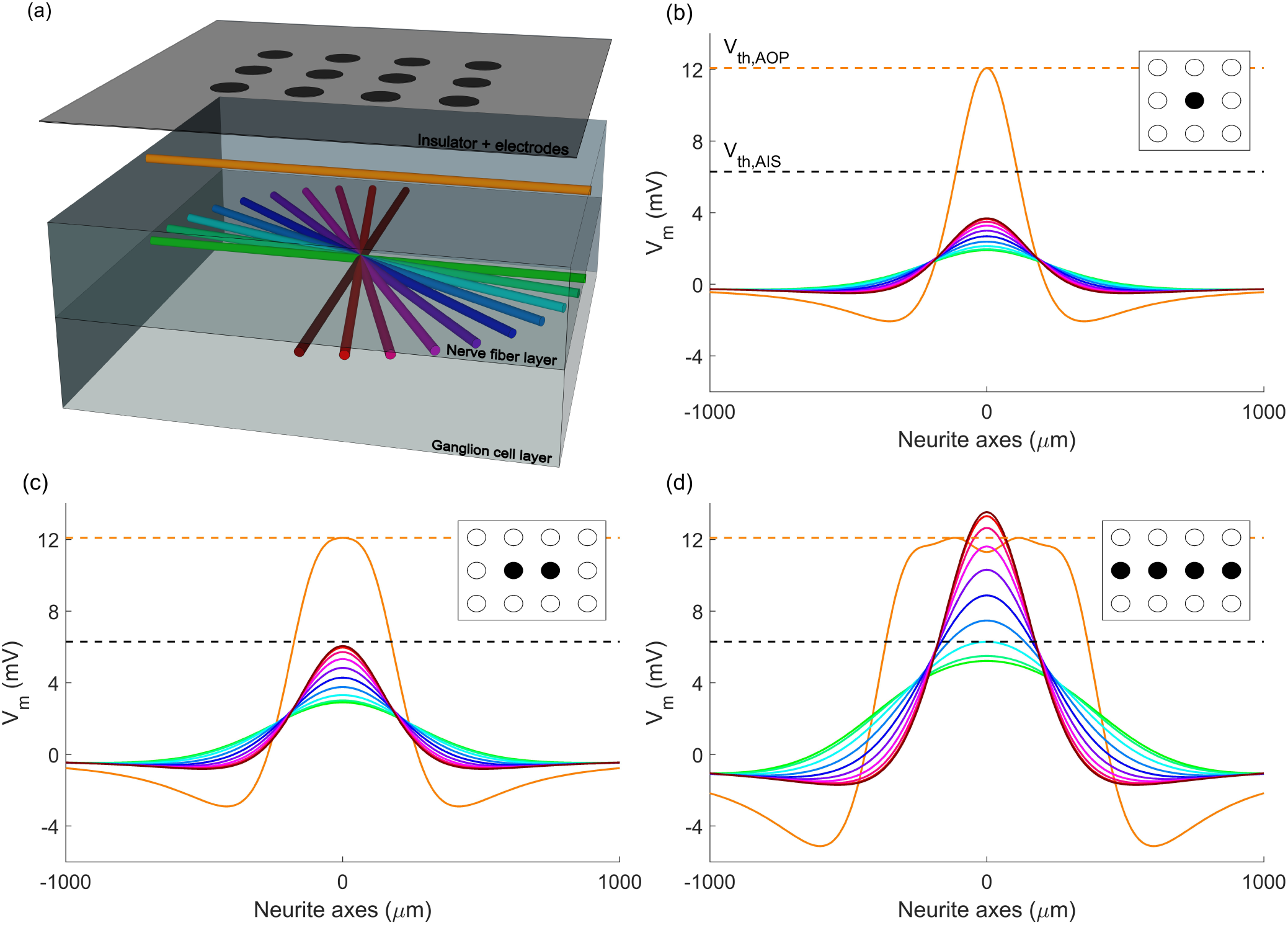
Geometry and simulated membrane potentials for axons of passage (AOPs) and axon initial segments (AISs) at a variety of *x-y* orientations. (a) Four layer model geometry showing the electrode array, an example of a parallel axon of passage (orange), and the neurite orientations considered in the ganglion cell layer (green-brown). Membrane potential at the end of the cathodic phase is shown along the axes of the neurites being simulated for configurations of (b) one, (c) two, and (d) four electrodes. Dotted lines represent membrane thresholds for axons of passage (orange) and axon initial segments (black). Stimulus currents have been chosen such that they drive the axon of passage precisely to its threshold level: 201.2*μ*A for (b), 342.5*μ*A for (c), and 870*μ*A for (d). Colors in (b)-(d) indicate corresponding neurites in (a).

Figs. 5(c) and 5(d) provide an initial assessment of the combined influence of tissue anisotropy and electric field orientation on activation of AOPs and AISs. The work of Rattay and Resatz [11] indicated that the activation of a passing fiber may be limited by controlling the way in which the induced electric field changes along the length of that fiber. Hence, simulations have been designed that recruit a number of electrodes aligned with the direction of the considered AoP. As can be seen from Fig. 5, the level of AIS versus AoP activation increases markedly as the number of electrodes increases. With four electrodes, it is possible to activate 78% of AIS fibers before activation of the overlying layer. When compared to Fig. 5(b), there is a consistent increase in the relative activation of perpendicular (green) and parallel (brown) AISs in the GCL for two and four electrode configurations. For comparison, the ratio of perpendicular to parallel AIS activation is 1.9, 2.1, and 2.5 for one, two, and four electrodes, respectively.

### C. Effect of pulse duration and electrode-retina separation

Using the threshold values determined above, a parameter sweep was conducted over pulse frequencies and electrode-retina separations. For each set of parameters, simulations were run to compare the membrane activation of parallel neurites in the NFL and neurites with a range of rotated orientations in the GCL. For both the single orientation in the NFL and the range of simulated fiber orientations in the GCL, membrane potential was calculated for fibers across the full plane at the appropriate retinal depth. Under the assumption that the orientation of axon initial segments is described by a uniform distribution, the proportion of preferentially activated AIS fiber orientations was determined. This is illustrated in Fig. 6, which shows the level of preferential activation achieved for a variety of stimulation parameter combinations. In this analysis, preferential activation is defined as when the membrane potential of an AIS is driven to its threshold potential at a lower stimulus current than is required to drive *any* AOP to threshold.

**Fig. 6.**
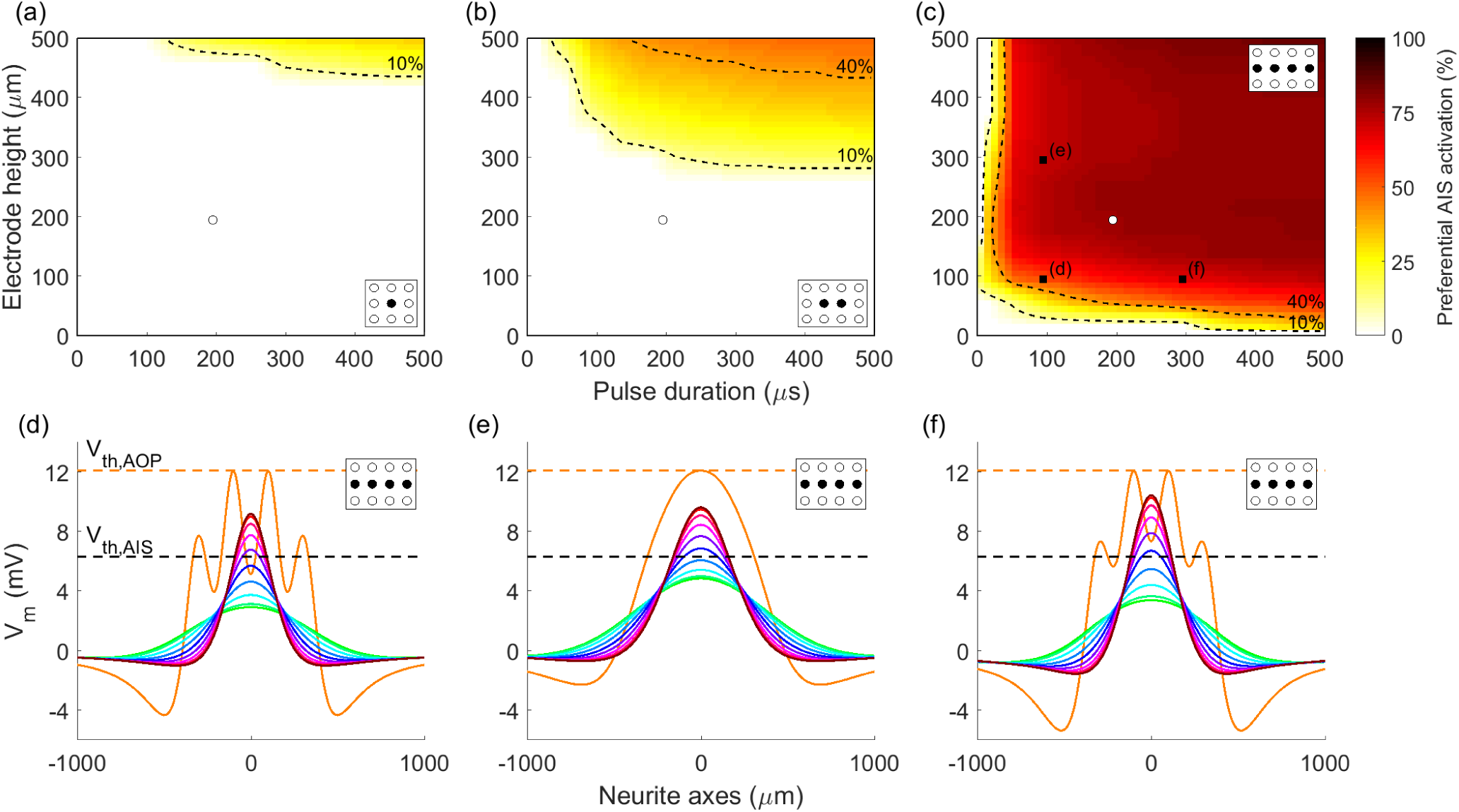
Proportion of AIS orientations preferentially activated for different electrode-retina separations (*d_ER_*) and pulse durations. Heat maps indicate the proportion of AISs activated at a lower stimulus current than any fibers in the NFL for (a) one-, (b) two- and (c) four-electrode configurations. Regions of low (<10%), medium (10–40%), and high (>40%) stimulation selectivity are separated by dotted contours. White markers indicate the parameters used in Fig. 5, and black markers indicate the parameters used for subplots (d), (e) and (f), which show examples of simulated membrane potentials for axons of passage and axon initial segments. Colors in (d)-(f) correspond to those in Fig. 5(a).

Fig. 6(a) highlights the challenge of acheiving preferential activation of the GCL using classical, single-electrode stimulation. Only very small levels of selectivity are obtained even with the most favourable stimulation parameters (large electrode height and pulse duration). A dramatic increase can be seen in the range of stimulation parameters at which preferential activation is achieved when moving from the one or two electrode configurations to four electrodes.

A comparison of the membrane potential induced by four-electrode stimulation with small and large elecrode-retina separation can be seen by comparing Fig. 6(d) and Fig. 6(e). A clear effect is that, due to the smoothing effect of increased current spread with greater electrode-retina separation, the AOP membrane potential along the axon has a much smoother shape for electrodes positioned further from the retina. Less obviously, larger separation distances result in increases in preferential activation of the GCL, due to the increased opportunity for summation of currents originating from adjacent electrodes. Similarly, as can be seen in Fig. 6(d) and Fig. 6(f), increases in pulse duration also result in increases in preferential activation of the GCL. Importantly, however, the overwhelming majority of the change in preferential activation occurs for pulse durations of less than 100*μ*s and separation distances of less than 100*μ*m.

### D. Performance of simultaneous four-electrode stimulation

An important assessment of these results is how the increase in preferential activation of the GCL affects key clinical performance metrics, such as the required total stimulus current and the spatial selectivity of activation, which is measured here using activation radius. In the following analysis, GCL activation level is defined as the percentage of AIS orientations that are activated (depolarised above membrane threshold) given a specific stimulus. This percentage is taken at the point in the plane of analysis that is maximally activated, which in all simulated examples is centered with respect to the electrodes. Activation radius is used to show the width of the region that is activated by a given stimulus, which will directly affect the resolution achievable with an implanted device and is defined as the radius of the smallest circle that encloses all areas with nonzero activation.

Fig. 7(a) shows the relationship between stimulus current and GCL activation level, and how this relationship changes with electrode configuration and electrode-retina separation distance, d_ER_. As expected, to achieve an equal level of activation for more distant electrodes, greater stimulus current is required. Fig. 7(b) shows the variation in activation radius with total stimulus current and Fig. 7(c) shows the correspondence between activation level and activation radius in the GCL. In each of Figs. 7(a)-(c), dashed curve regions indicate configurations in which AOPs are activated preferentially to AISs. In terms of isolating the optimal stimulus level, it is important to consider whether or not this will result in co-activation of passing axons as indicated by dashed regions, the level of activation achieved in the GCL, and the resulting radius of activation in the GCL. To facilitate comparison of the spread of activation in the GCL induced by one-, two-, and four-electrode configurations, two-dimensional maps of activation in the *x-y* plane are shown in Figs. 7(d)-(f), along with the locations of the stimulating electrodes. Importantly, despite utilising four times more electrodes, the activation radius at a given activation level for the four-electrode configuration is typically less than 200% of the activation radius for one electrode, as indicated by dotted lines in Fig. 7(c). Furthermore, very little additional total current is required for four electrodes compared to one electrode, so that current per electrode is considerably less than for single electrode stimulation.

**Fig. 7.**
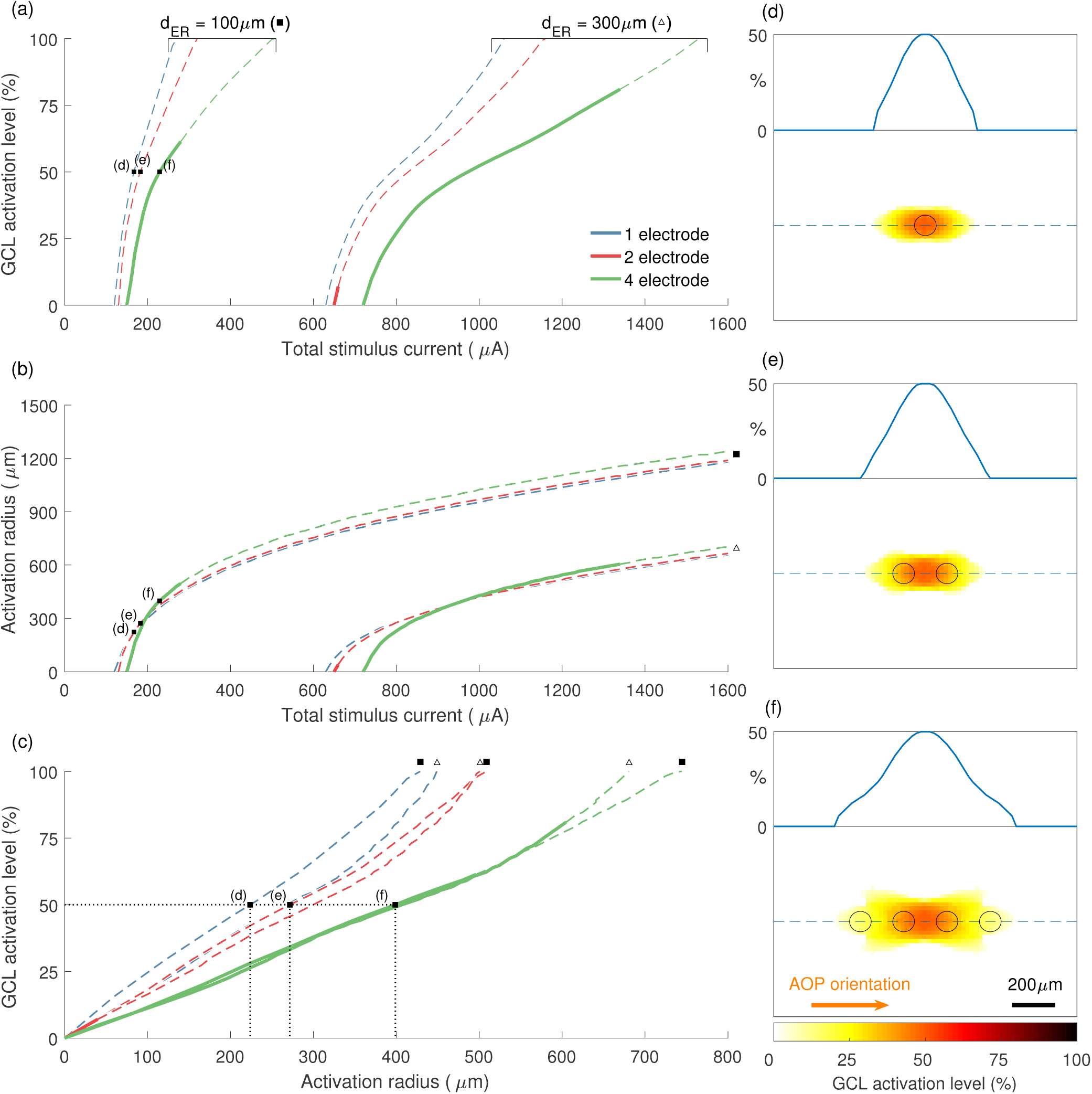
Performance of different electrode configurations with respect to GCL activation level, required stimulus current, and radius of activation. (a) The proportion of AIS orientations activated vs. total stimulus current for various electrode configurations and electrode-retina separation distances, dER. (b) The radius of the activated region vs. total stimulus current. (c) The relationship between activation radius and activation level. Stimulation strategies analysed in (a)-(c) include one-, two-, and four-electrode configurations, as well as separation distances of 100μm (filled square) and 300*μ*m (unfilled triangle). Solid and dashed regions in (a)-(c) represent configurations that result in preferential activation of AISs and preferential activation of AOPs, respectively. Labelled points in (a)-(c) correspond to the examples plotted in (d)-(f), which show the spread of GCL activation in the *x-y* plane. Dashed blue lines in (d)-(f) correspond to one-dimensional insets. All simulations used a pulse phase duration of 200*μ*s.

### E. Non-ideal electrode array placement

In reality, electrodes are unlikely to be aligned with passing axons, as in Fig. 7(f). This is due to both the placement of the implanted device and the curvature of passing axons as they pass under the electrode array. In order to test the application of the multi-electrode stimulation strategy for non-ideal electrode placement, several more challenging geometries were simulated. In each case, the electrodes recruited for stimulation were chosen to represent the most logical extension of the ideal four-electrode configuration presented above, and the electrodes were stimulated with equal current.

Fig. 8 shows an assessment of two such geometries: one where the target for stimulation is centered between four electrodes and another where the target for stimulation is centered between two electrodes and with a non-parallel axon of passage orientation of 22.5 degrees, as shown in the insets in Figs. 8(a)-(b). For the former case, stimulation current was delivered by eight electrodes in total. Another obvious choice of AOP orientation to analyse is 45 degrees. However, because this orientation aligns with diagonal rows of electrodes, the outcome was very similar to the ideal, zero degree case and so has been omitted here.

**Fig. 8.**
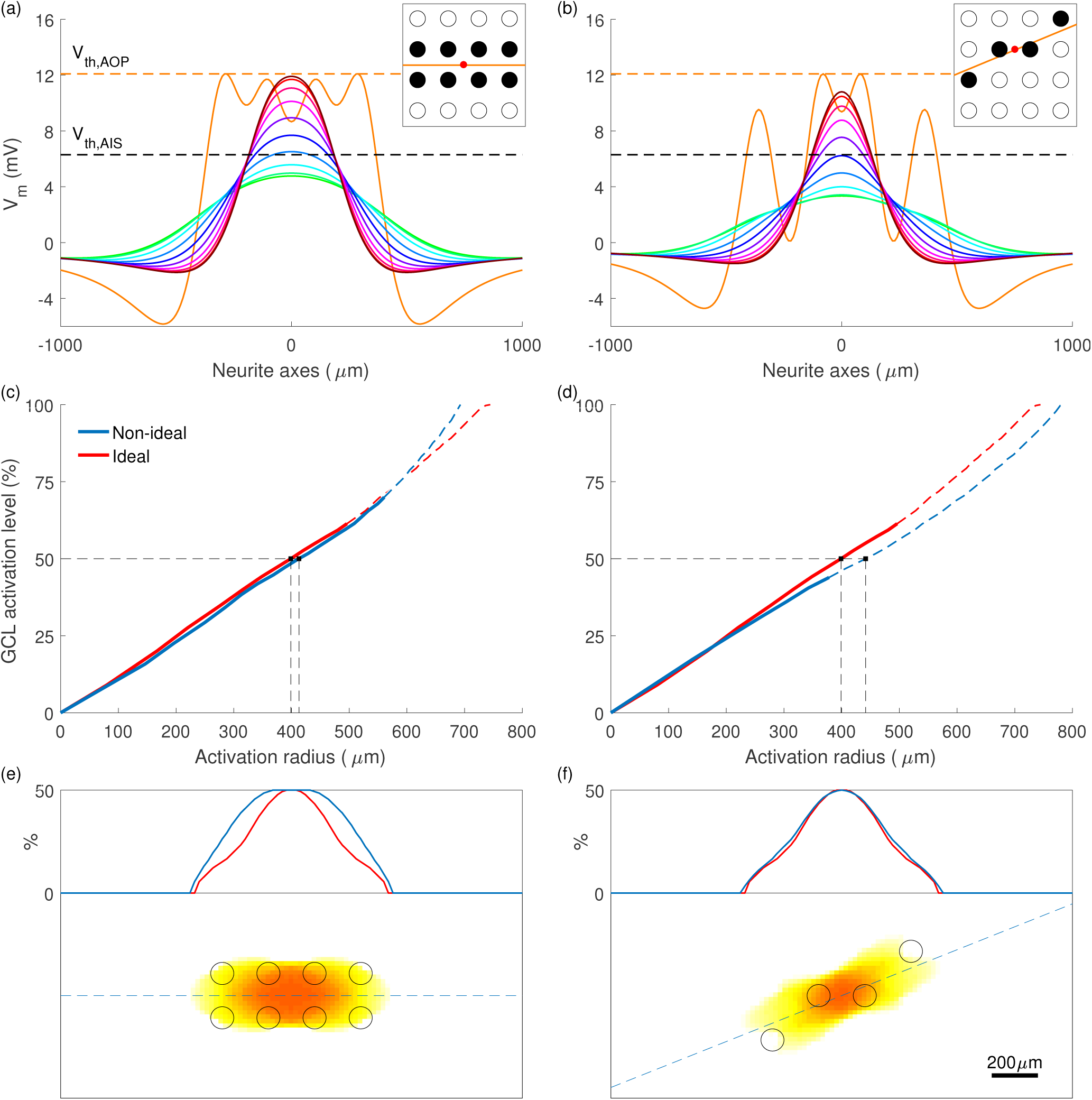
Preferential stimulation for two non-ideal electrode array placements. (a)-(b) Membrane potential along neurite axes for axons of passage and axon initial segments, with stimulus current chosen to maximally activate initial segments without activating any passing axons. Colors correspond to those in Fig. 5(a), with green parallel to axons of passage and brown perpendicular. Insets describe the geometry of each simulation, indicating target region (red), electrodes used (black), and the orientation of axons of passage (orange). (c)-(d) GCL activation level vs. activation radius for non-ideal and ideal (as in Fig. 7(f)) geometries. Transitions from solid to dashed lines represent the transitions from axon initial segment to axon of passage preferential activation. (e)-(f) The spread of GCL activation in the *x-y* plane. The dashed blue line corresponds to the one-dimensional inset. All simulations used a pulse phase duration of 200*μ*s and electrode-retina separation of 100*μ*m.

The resulting membrane potential along the axis of an AOP and AISs with varied orientations is presented in Figs. 8(a)-(b). For each configuration, preferential activation of AISs was achieved, with 70% and 44% of AIS orientations being activated at lower stimulus currents than any axons of passage for the eight- and four-electrode configurations shown, respectively (compared to 61% for the ideal four-electrode configuration). As shown in Figs. 8(c)-(d), the relationship between GCL activation and activation radius is comparable with that of the ideal configuration. Finally, the *x-y* activation maps in Figs. 8(e)-(f) indicate only modest increases in the spread of activation when compared to the ideal case.

## IV. Discussion

### A. Key factors influencing preferential retinal activation

The observed dependence of activation on neurite orientation is a result of several competing factors. The dominant orientation of axons in the NFL results in highly anisotropic spread of extracellular potential under stimulation. As a result, the orientation of fibers in the GCL with respect to this anisotropy influences membrane potential. The overall probability of eliciting a response selectively in the GCL then depends on the relative influence of fiber rotation, membrane threshold, and fiber depth.

Fig. 9 shows the change in the spread of extracellular potential in directions parallel and perpendicular to the orientation of AOPs. Current spreads through the NFL much more readily in the direction of the overlying fiber tracts than it does perpendicularly to them. This leads to a more rapid change in extracellular potential when moving away from stimulating electrodes in the direction perpendicular to the AOPs. This, in turn, results in the directional spatial derivatives of extracellular potential being greater in this perpendicular direction, leading to maximal activation of AISs with perpendicular orientation in the GCL, as seen in Fig. 5(b). Specifically, the activation due to orientation is influenced via differences in the second spatial derivative of the extracellular potential, which manifests in the frequency domain in Equation (16a) as 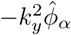. This analysis also shows that modulating the spread and orientation of the electric field by driving multiple, aligned stimulating electrodes can be used to minimize activation of fibers with specific orientations, such as passing axons.

**Fig. 9.**
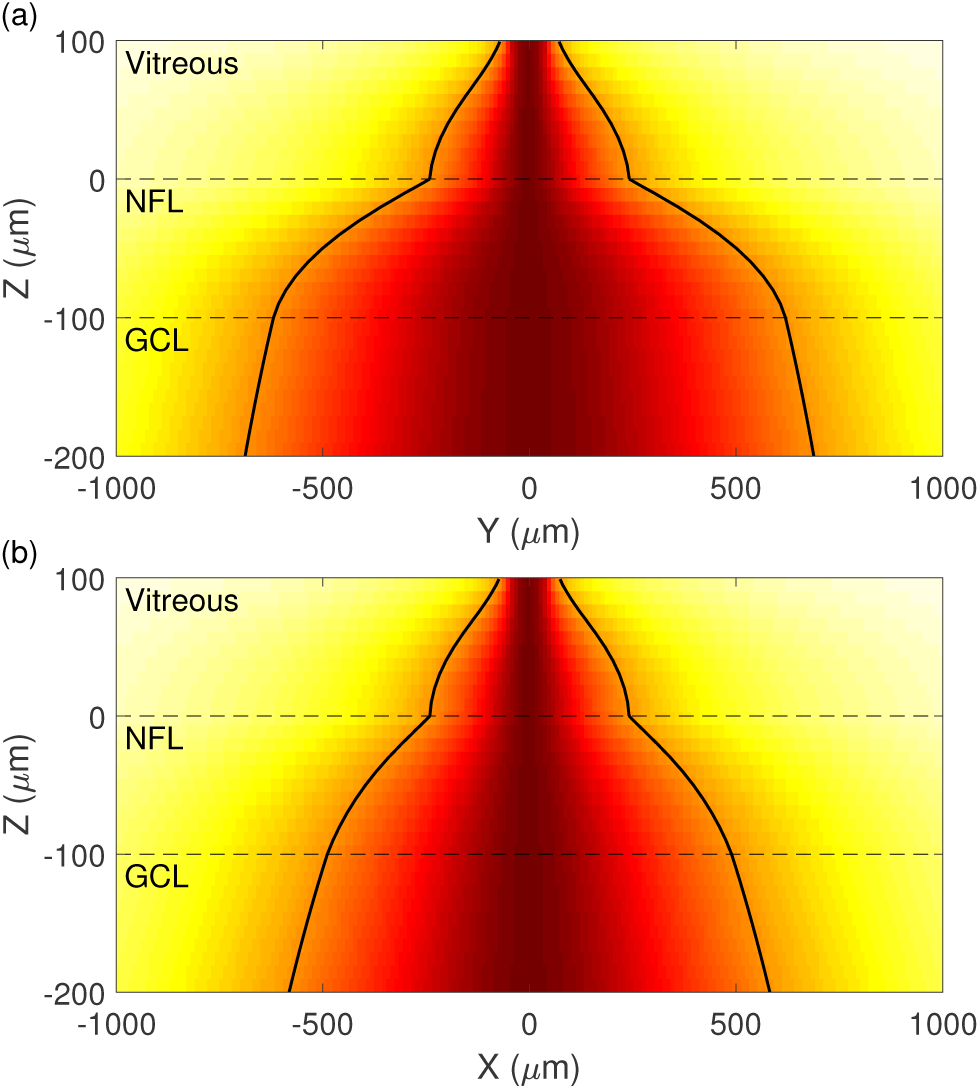
Normalized spread of extracellular potential with distance from a stimulating electrode. Spread is shown in (a) the *y-z* plane, parallel to the orientation of AOPs, and in (b) the *x-z* plane, perpendicular to the orientation of AOPs. The simulated extracellular potential at each z-slice is normalized to the range [0, 1] by subtracting the minimum and scaling the maximum per slice to 1. This is done for illustrative purposes due to the rapid fall-off of extracellular potential with increasing distance from the electrode. Contour lines indicate the half-width at full-maximum potential. Stimulation is with a single electrode located 100*μ*m above the retinal surface at the origin in the *x-y* plane. Dashed lines indicate layer boundaries.

A side effect of using multiple electrodes aligned with passing axons is that the ratio of depolarisation of perpendicular AISs to parallel AISs increases with the number of electrodes. In the results summarised in Fig. 5, the ratio of the maximal depolarisation for perpendicular AISs to parallel AISs is 1.9, 2.1, and 2.5 for one-, two-, and four-electrode configurations, respectively. The cause of the increase from one to four electrodes is likely that by aligning electrodes with passing axons, the activity of similarly oriented AISs in the GCL is also reduced, while having little effect on the depolarisation of perpendicular AISs. This effect is far less pronounced for parallel fibers in the GCL when compared to the NFL due to the natural spread of current at greater retinal depths; the artificial spread of current introduced by using multiple electrodes is less pronounced when compared to the natural longitudinal spread caused by the geometry of the NFL. In contrast, the anisotropic spread introduced by the NFL, shown in Fig. 9, has little effect on superficial AOPs as they are close to the retinal surface where current spread is still predominantly isotropic and so we must rely on electrode configuration to control the profile of extracellular potential.

A related phenomenon is highlighted by Fig. 7(c), which shows that, given a certain level of GCL activation, there is an increase in the spread of activation as electrodes are moved further from the retinal surface; however, this effect is not seen for four electrodes. The reason for this is that, due to the wider distribution of current at the electrode array, the increase in spread due to greater electrode-retina distance is marginal. Another key feature of the system is that the region in which the largest spread occurs is in the NFL in the direction of passing axons. Therefore, an increase in the distance of vitreous fluid through which current flows has a less pronounced effect on total spread in that direction. An increase in the spread of activation in the direction perpendicular to passing axons can be observed as electrode are moved away from the retina, however spread is always more pronounced in the direction of passing fibers.

Although these results are based on simulations of cylindrical neurites, the developed method for the analysis of arbitrarily rotated fibers can be applied directly to the simulation of unbranched axons with arbitrary morphology, as discussed in Section II-F. A preliminary next step will be to validate the current results using ganglion cell axon reconstructions. A key point of interest will be whether or not the effect is maintained when axonal orientation changes along the length of the simulated fiber. This will depend on the length constants associated with both axon curvature and membrane activation. If the latter is relatively smaller, axon curvature will have little effect and localized fiber orientation will determine the level of activation along the axon.

### B. Choosing a stimulation strategy

As can be appreciated from Figs. 6(a)-(c), of the electrode configurations that were simulated, preferential stimulation with clinically desirable parameters can only be achieved with four electrodes. Ideally, electrodes should be placed as close as possible to the surface of the retina without causing damage. This reduces the required stimulus current and limits current spread, thereby increasing the achievable device resolution. From Fig. 6(c), the majority of the change in AIS activation with varying electrode height is seen to occur in the first 100*μ*m, suggesting that the optimal electrode height considering both preferential AIS activation and activation radius is around 100*μ*m. Beyond this height, little is gained in terms of preferential activation, with reductions in resolution and larger required stimulus currents.

Given the electrode-electrode separation used in this study, for separation distances of less than 50μm, preferential stimulation is limited due to a lack of lateral summation of currents from adjacent electrodes; activation under each electrode will occur in a similar way to one electrode. This highlights the fact that these results rely on current spread from adjacent electrodes overlapping and summing together. The level of this summation depends on both the distance between electrodes (the *x-y* distance that current has to spread) and the distance from the electrodes to the retinal surface (the *z* distance over which current can spread). In theory it is expected that, in the limit of infinitesimally small electrodes that are infinitesimally close together, preferential activation could be achieved with electrodes arbitrarily close to the retinal surface. In reality, the optimal electrode-retina separation distance will depend on the geometry of the electrode array and may differ from the results presented here.

The combination of Figs. 6 and 7 provide a starting point for choosing a clinically relevant stimulation strategy. Assuming that the height of the electrode array above the retina is fixed at 100μm and pulse duration is greater than 50μs, the chosen pulse duration has little influence on activation provided appropriate current magnitudes are delivered. Key remaining considerations are the required current, level of activation in the GCL, and size of the activated region, which can be determined from Fig. 7. It is unclear exactly how either GCL activation level or activation radius in the current model will map to perception by patients with an implanted device. As such, a suitable stimulus current may need to be determined either experimentally or based on direct feedback from device users. This current level will depend on the trade-off between GCL activation level and activation radius (Fig. 7(c)), and should always be kept below the level required for AOP-related perception, and within clinically determined safety limits. As an example, for four-electrode stimulation with an array positioned 100μm above the retina, this current level is 280μA, as indicated by the transition from solid to dotted lines in Figs. 7(a) or (b). A valuable implication of using four-electrode stimulation is that it requires only small increases in total stimulus current to achieve a similar level of activation in the GCL when compared to single-electrode stimulation. This results in a 3–4 times decrease in the required current per electrode or per area, which is the most clinically relevant measure safe stimulation current.

Although Fig. 7 shows that by recruiting more stimulating electrodes the induced area activated becomes greater, it should be noted that this will not necessarily reduce perceived resolution. Previously, recipients of epiretinal implants have reported elongated and line-like phosphenes, thought to be caused by stimulation of passing axons in the NFL that originate from distant regions of the GCL [1, 11–19]. Hence, despite an increase in the region of activation in the GCL when using a four-electrode stimulation strategy, the overall resolution is expected to increase due to the elimination of activation of the NFL. Furthermore, phosphene regularity is expected to be greater under the proposed strategy, more readily facilitating the development of more complex stimulus patterns built up from this perceptual subunit.

### C. Determining membrane thresholds

To our knowledge, although stimulus current thresholds have been reported for the AIS and distal axon of RGCs, there exists no experimental data on the membrane thresholds of RGCs at these locations. In previous models, the low threshold of the AIS has only been incorporated into active, conductance-based models of RGCs. In these models, the threshold is reduced at the AIS by increasing the sodium channel density by a factor ranging from 2 to 40, generally chosen to reproduce desired physiological responses [15, 19, 51].

In order to choose appropriate thresholds for our passive model and to avoid arbitrarily choosing a reduced threshold, membrane threshold levels were determined using an approximate reproduction of the experimental procedure of Fried et al. [42]. Simulating the experiment using the same modeling framework in which the thresholds were later applied ensured that the chosen values were representative of the reported experimental data and were relevant to the current model. The ratio of the calculated membrane threshold for the AOP and AIS was approximately 2, representing the lower end of the ratios used in other models, and suggesting that the current outcomes are conservative. It was assumed that, although approximating the conical electrodes used experimentally by disc electrodes may change the determined threshold values slightly, it was unlikely to have a large effect or to alter their ratio.

### D. Experimental validation

Controlled experimental validation of these results requires techniques for the measurement of RGC activation at multiple locations in the retina simultaneously. *In vitro* studies in which the average trajectory of passing axons in the NFL is known will allow for measurements of activation to be taken in the GCL at both the region being targeted by stimulation and at more distant locations that lie under the trajectory of passing axons. Methods have also been developed for imaging GCL activity across the whole retina [17]. A challenge with quantitatively validating the result in this paper is that the small distance between the electrode array and the surface of the retina must be very tightly controlled.

Due to the dependence of these results on the anisotropy of the NFL, it is expected that varying the thickness of the NFL will have a marked effect. The chosen NFL layer thickness is based on an approximation of the human retina, and so these results are relevant only to human retinal stimulation. Rodent models used for research and testing of epiretinal implants have thinner NFLs, and so the influence of retinal layer orientation will be less pronounced. Although this in no way confounds the current findings, it suggests that experimental validation would be best carried out in the primate retina. A potential solution for other animal models may be to modify the present model to represent the appropriate animal model so that any observed evidence can be extrapolated to human-like retinal geometries. It is important to note that a large part of the present result derives from electrode configuration, which can be kept consistent across different animal models.

### E. Optimizing electrode currents

For stimulation strategies that utilize more than two electrodes, it is likely that delivering equal currents to all electrodes does not represent the optimal stimulus for achieving preferential activation with minimal activation radius. It may be possible to more optimally distribute currents across the recruited electrodes in a way that minimizes the activating function along AOPs. As can be seen from Figs. 7(d)-(f), for one- and two-electrode configurations, the profile of activation about the centre of the electrode array follows a simple curve with monotonic first derivative. In contrast, with four electrodes, the profile has a more complex shape due to the added degree of freedom. In this case, this extra degree of freedom can be represented by the ratio of the current delivered to the two internal and two external electrodes.

As highlighted by Fig. 8, there is a range of electrode/AOP orientations that must be dealt with by a proposed stimulation strategy. We have demonstrated that the approach proposed in this paper is robust to changes in relative electrode array to AOP orientation, and can target off-centered tissue volumes. However, it is again likely that delivering equal currents to each electrode is sub-optimal. In this more general case, optimal electrode currents will also depend on the particular pattern of electrodes that is being used. For instance, the optimal ratio of internal electrode currents to external electrode currents for the case presented in Fig. 8(b) will be different than for a set of four electrodes perfectly aligned with the AOP.

With four-electrode stimulation, which can be tuned by a single parameter, optimization could be achieved using a simple brute force search through possible current ratios. However, the model presented here is linear and has an analytic solution in the Fourier domain. This means that a closed-form solution to this optimisation problem can be found using least squares or some other linear optimization algorithm. This approach could be applied to the optimization of currents delivered to an arbitrary number of electrodes in order to minimise activation of the NFL. Optimization of multiple electrode currents to achieve both focal activation of the GCL and minimal activation of the NFL will be the subject of a subsequent study.

## V. Conclusion

This paper demonstrates that activation of RGCs in the inner retina under epiretinal stimulation depends on both axonal orientation and orientation of the stimulating electric field relative to the orientation of axons of passage in the NFL. The developed model allows for an analysis of this dependence by capturing the distinct distributions of fiber orientation of the nerve fiber layer and the ganglion cell layer. A four-electrode stimulation strategy has been proposed that accomplishes preferential activation of the retinal ganglion cell axon initial segment over passing axons in the NFL using clinically suitable stimulus currents and electrode configurations. Although concessions must be made with regard to activation radius in the GCL, these are relatively minor, and the proposed strategy is expected to enable higher resolutions and more clearly interpretable percepts by users of epiretinal prostheses.

## VI. Acknowledgements

TE was supported by an Australian Postgraduate Award from the Australian Government and The University of Melbourne, the Gowrie Scholarship Fund of the Australian National University, and IBM Research, Melbourne. RRK acknowledges the support of IBM Research, Melbourne where he was employed for a portion of this research. HM acknowledges funding from the Australian Research Council Centre of Excellence for Integrative Brain Function (project number CE140100007). ANB acknowledges the support of the Australian Research Council’s Discovery Projects funding scheme (project n<u>um</u>ber DP140104533). DBG, HM, and ANB acknowledge the support of the Australian National Health and Medical Research Council’s Project Grant funding scheme (NHMRC Grant APP1106390). This research was supported by the Victorian Life Science Computation Initiative (VLSCI) on its Peak Computing Facility at The University of Melbourne (grant number VR0138). Kerry Halupka, Ewan Nurse, and Philippa Karoly are thanked for providing feedback and discussion.

